# Short-Term Evolution and Dispersal Patterns of Fluconazole-Resistance in *Candida auris* clade III

**DOI:** 10.1101/2024.05.22.595305

**Authors:** Irving Cancino-Muñoz, Juan Vicente Mulet-Bayona, Carme Salvador-García, Nuria Tormo-Palop, Remedios Guna, Concepción Gimeno-Cardona, Fernando González-Candelas

**Affiliations:** Unidad Mixta Infección y Salud Pública FISABIO-Universidad de Valencia, Valencia, España; Instituto de Biología Integrativa de Sistemas, I2SysBio (CSIC-UV), Valencia, España; Servicio de Microbiología y Parasitología, Consorcio Hospital General Universitario de Valencia, Valencia, España; Departamento de Microbiología y Ecología, Universidad de Valencia, Valencia, España; CIBER en Epidemiología y Salud Pública, ISCIII, Madrid, España

**Keywords:** Fungal pathogen, *Candida auris*, fluconazole resistance, phylogenetic structure, dispersion

## Abstract

The rapid increase of infections caused by the emerging fungal pathogen *Candida auris* is of global concern, and understanding its expansion is a priority. The phylogenetic diversity of the yeast is clustered in five major clades, among which clade III is particularly relevant, as most of its strains exhibit resistance to fluconazole, reducing the therapeutic alternatives and provoking outbreaks that are difficult to control. In this study, we have investigated the phylogenetic structure of clade III by analyzing a global collection of 566 genomes. We have identified three subgroups within clade III, among which two are genetically most closely related. Moreover, we have estimated the evolutionary rate of clade III to be 2.25e-7 s/s/y (2.87 changes per year). We found that one of these subgroups shows intrinsic resistance to fluconazole and is responsible for the majority of cases within this clade globally. We inferred that this subgroup may have originated around December 2010 (95% HPD: April 2010 - June 2011), and since then it has spread across continents, generating multiple large outbreaks, each with a unique pattern of transmission and dissemination. These results highlight the remarkable ability of the pathogen to adapt to its environment and its rapid global spread, underscoring the urgent need to effectively address this epidemiological challenge.

**IMPORTANCE:** The number of cases affected by *Candida auris* has increased worryingly worldwide. Among the currently recognized clades, clade III has the highest proportion of fluconazole-resistant cases and is spreading very rapidly, causing large nosocomial outbreaks across the globe. By analyzing complete fungal genomes from around the world, we have confirmed the origin of this clade and unraveled its dispersal patterns in the early 2010s. This finding provides knowledge that may be helpful to the public health authorities for the control of the disease.

## INTRODUCTION

Infections caused by the emerging pathogen *Candida auris* have increased dramatically worldwide (1–3). This fungus is characterized by causing large, difficult-to-control nosocomial outbreaks (4–6), as well as by its rapid acquisition of resistance to antifungals (1, 7). As a result of this epidemiological situation, the World Health Organization has recently included it in its fungal priority pathogens list (8).

The phylogenetic structure of *C. auris* consists of 5 major clades with different genomic and phenotypic characteristics. For instance, clade I, originated in South Asia, is characterized by its rapid dissemination in clinical settings and is responsible for outbreaks that are difficult to eradicate worldwide (9–11). In contrast, strains belonging to clade V have only been detected in individuals from Iran (12). Most of the strains classified in clade III are resistant to fluconazole, an antifungal agent used as a first-line treatment against *Candida* infections (13).

Global phylodynamic studies suggest that the oldest clade is clade II, while the most recent ones are clades I and III (4). Estimates of the mutation rate of *C. auris* vary between 1.34e-5 - 5.74e-5 substitutions per site per year (s/s/y), depending on the clade analyzed and the epidemiological context of each study (4, 14, 15).

Since 2016, the number of *C. auris*-associated infections in Europe has tripled, with the Mediterranean countries being the most affected. In the latest ECDC report, Spain, Italy and Greece were responsible for 96.3% of the cases in this continent. Of these, Spain is the largest contributor with 50.5% of the total (16). National genomic surveillance studies in Italy have identified the emergence of multidrug-resistant strains in a large hospital in Liguria since 2019, all belonging to clade I (9). The situation in Spain is similar. Monitoring studies in different medical centers have detected *C. auris* since 2016. In all cases, the strains have been classified into clade III, and were resistant to fluconazole, but only a few have been denoted as multidrug-resistant (6, 17).

The origin of clade III is thought to be South Africa. There have been cases reported in 2011 and 2012 in Kenya and South Africa (4, 18). Nevertheless, there are reports describing cases belonging to this clade in Canada in 2012, so the origin of this clade remains controversial (19). Genomic epidemiology studies of nosocomial outbreaks in other continents have shown that the strains causing these outbreaks were genetically more closely related to African than to North American strains.

In this study, we have analyzed the evolution and dispersal of clade III of *C. auris*, focusing on two different epidemiological contexts, one regional, analyzing the initial stages of a long-term nosocomial outbreak in Spain, and the other global. Using information from a large collection of genomes, we describe three genetically distinct subgroups within clade III, one of which is intrinsically resistant to fluconazole. The mutation rate obtained as well as the most likely inferred ancestral dates provide new insights into understanding how this clade has evolved and novel information for the genomic surveillance of this emerging pathogen.

## MATERIALS AND METHODS

### Sample collection and whole genome sequencing

This study focuses on the evolutionary analysis of the global dispersion of clade III of *C. auris.* Hence, the analyzed dataset is mainly enriched with samples from this clade; nevertheless, we included samples from the other *C. auris* clades to determine the genotype of the unknown isolates. The search criteria for public samples in the NCBI database were as follows (latest access on 2023/06/14): records from clinical and/or environmental isolates, generated with short-read sequencing technology, coming from all continents, and containing "clade III" in their genotype description, when possible. Only records containing at least the year of diagnosis in the variable "collection date" were selected. Importantly, all the samples belonging to three previously characterized clade III nosocomial clusters in the UK (20), China (5), and Kenya (4) were downloaded. Additionally, we included 35 representative samples of a nosocomial outbreak in the Valencia Region, Spain, one of which was sampled in the hospital environment (21). In cases with more than one sample per patient (e.g., serial samples from the same individual), we selected the first diagnosed sample from each case, when possible. Records with less than a minimum depth of 20x and minimum horizontal coverage of 90% were discarded (n=17). Thus, our final dataset consisted of 689 *C. auris* genomes (clade III: n=567; other clades: n=122).

Genomic libraries of isolates from the Valencian region outbreak were prepared using the Illumina DNA Prep kit (Illumina, San Diego, CA, USA) and sequenced on a Nextseq instrument generating 2×150bp paired-end sequencing reads following the manufacturer’s instructions.

### Bioinformatic analysis

All raw sequencing reads were trimmed and filtered using fastp (22) with the following parameters “--cut_tail, --cut_window_size=10, --cut_mean_quality=20, -- length_required=50, --correction, and --dedup”. These filtered reads were used to detect mutations associated with antifungal resistance as well as for phylogenetic analyses as described below. In the case of the reference strains, we first downloaded their respective assembly file, and then short-reads were simulated using the ART software (23) with the following parameters “-l 150 -ss MSv3 -f 100 -m 350 -s 10”. The reference assembly samples were “GCA_002759435.2” (clade I), “GCA_003013715.2” (clade II), “GCF_002775015.1” (clade III), and “GCA_003014415.1” (clade IV).

The Valencian region raw sequence reads were deposited in the European Nucleotide Archive public server under project PRJEB70513. Additional data and accession numbers of the sequenced reads can be found in **Supplementary Tables S1-S4**.

### Variant calling and identification of antifungal resistance mutations

We used two reference genomes for the identification of variants (indels and SNPs) depending on the goal of each analysis. The clade III reference genome (strain B12221, GCF_002775015.1) was used for evolutionary and phylogenetic analyses, whereas the fully susceptible clade I reference (strain B8441, GCA_003014415.1) was chosen for screening of mutations associated with antifungal resistance. In both approaches, the snippy v4.6.0 software (24) was employed with parameters "-- mincov 10 --minfrac 0.9 --minqual 60" for variant calling. For evolutionary analyses, polymorphisms detected in duplicated and repetitive regions were filtered out. Duplicated regions were identified with MUMmer 4.0.0 (25) while repeat regions were identified using RepeatModeler2 (26). In total, 3842 regions totaling 409,448bp (∼3.21% of the genome) were removed from posterior analyses. The presence of resistance-associated mutations in the *erg11* and *fks1* genes was determined by annotation of the variants with snpEFF (27).

### Phylogenetic and evolutionary analyses

Three different multiple sequence SNP-based alignments (MSAs) were constructed. An initial MSA was obtained including all global genomes (n=689) previously described, to place all the samples in a global phylogenetic context. The second MSA was generated using only the genomes belonging to clade III. Finally, a third MSA was constructed for the dating analysis, which included samples belonging to subgroups 2 and 3 of clade III (a detailed description of this analysis is given below).

All the alignments were reconstructed from the SNPs identified by mapping against the reference genome of clade III. In all the cases, the phylogenies were generated with Iqtree2 (28) using the GTR model with 1000 replicates and with the *-fconst* option added for each case. Finally, pairwise SNP distances were calculated for all MSAs with the R ape package.

### tMCRA inference and Bayesian skyline analysis

For the dating analysis, 380 global genomes with precise date of sampling (at least month and year) were used. This dataset included samples corresponding to four previously described nosocomial clusters, one from the UK, one from China, and one from Kenya, as well as a local outbreak (Valencia Region). First, root-to-tip distances were calculated with Tempest v1.5.3 (29) to determine whether sufficient temporal signal was present and to obtain an initial estimate of the mutation rate. Once we were confident that the temporal signal was strong, a Bayesian-based phylogeny was constructed using the collection date of each isolate in Beast2 (29) using a GTR nucleotide substitution model, with a Coalescent Exponential Population tree as a prior, and under a strict molecular clock with a LogNormal distribution prior. Furthermore, sampling bias correction was performed following the ascertainment correction recommendation (https://www.beast2.org/2019/07/18/ascertainment-correction.html). Three independent runs of 50 million MCMC chain lengths were conducted. The estimates obtained were inspected with Tracer (30). All the runs and trees were combined and annotated with LogCombiner and TreeAnnotator v2.7.1, respectively. The maximum credibility tree was visualized with FigTree v1.4.4.

## RESULTS

### Phylogenetic structure of *Candida auris* clade III

From a global public dataset containing 676 genomes, 22 genomes that did not meet our quality control criteria were filtered (see methods). In addition, 35 genomes belonging to an ongoing nosocomial outbreak in Valencia were added. For the reconstruction of the initial phylogenetic tree, we excluded polymorphisms detected in duplicated and repetitive genomic regions. In total, 3842 regions spanning 409,448 base pairs (∼3.21% of the total genome) were filtered out for further analysis. Thus, our initial dataset comprised 689 genomes and an alignment of 194,389 SNPs.

An initial ML phylogenetic tree was constructed using a global dataset that included isolates from all *C. auris* clades. We found that 566 genomes belonged to clade III (**Supplementary Figure S1**). Because most isolates belonging to this clade are fluconazole-resistant and cause other outbreaks that are difficult to control in Europe and globally, we decided to investigate this clade further.

In order to have a higher resolution of the phylogenetic structure of clade III, we inferred a ML phylogenetic tree using only the genomes from this clade of the global dataset (n=566, MSA consisted of 2,152 SNPs). The resulting topology showed that clade III is composed of 3 well-defined monophyletic subgroups (denoted as SG1, SG2, and SG3 in this study and corroborated by bootstrap support values of 100% in all cases) (**Figure 1**). Globally, subgroup SG3 is the most widespread, with 562 isolates from 9 countries in all the continents. The country with most isolates in SG3 was the USA with 279 isolates, followed by South Africa and the United Kingdom with 107 and 51 cases each, suggesting that this subgroup has spread worldwide. In contrast, subgroups SG1 and SG2 have only 2 isolates each. Those in the former were sampled in Canada, while those in SG2 are from South Africa (**Figure 1**). Examining the global phylogeny of clade III, we observed that isolates in subgroups SG2 and SG3 are the most closely related genetically, with mean genetic distances of 63 SNPs (ranging from 49-83 SNPs), while SG1 is the most distant with a mean genetic distance of 773 and 765 SNPs to SG2 and SG3, respectively.

**Figure 1.**
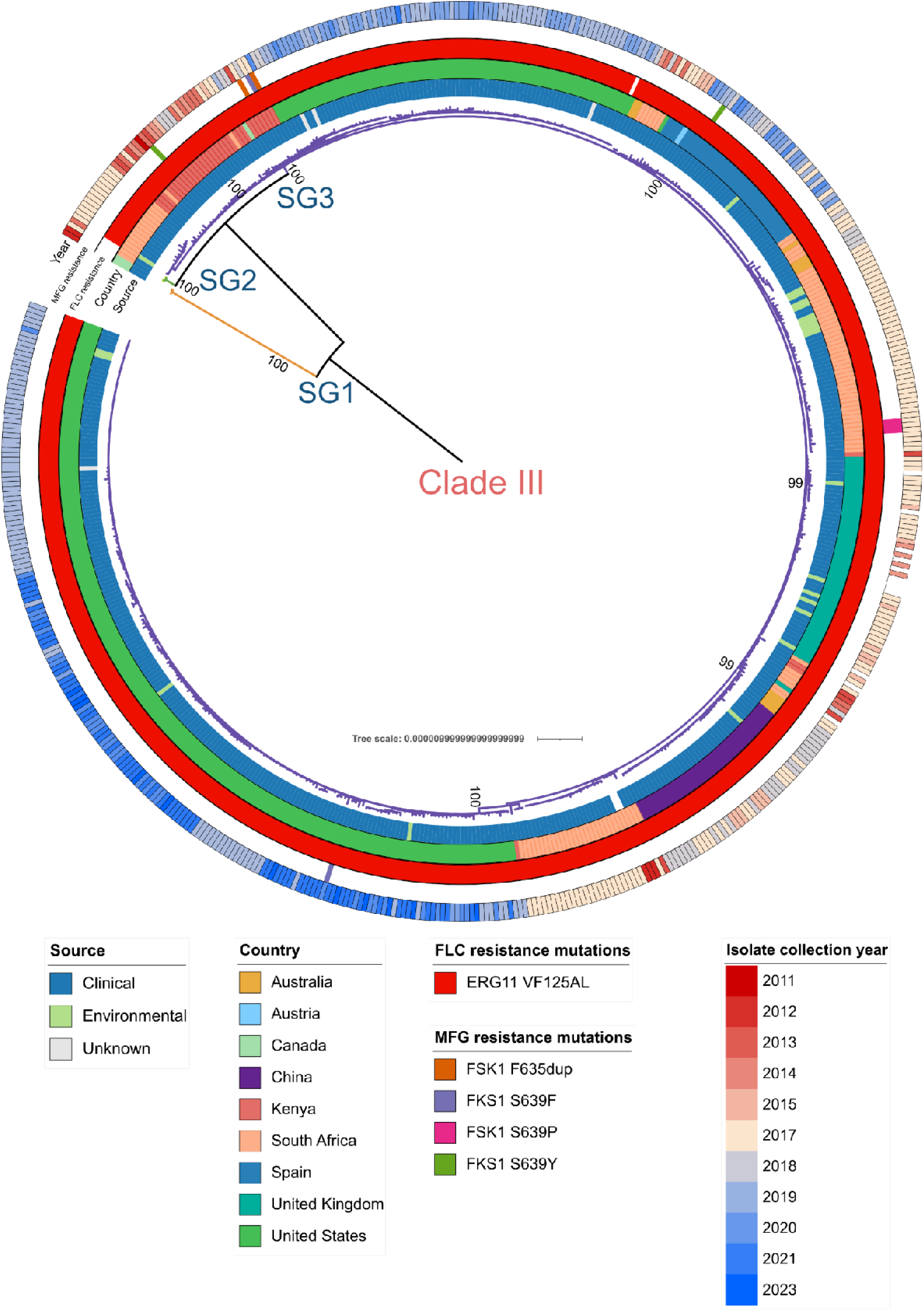
Phylogenetic structure of clade III of *Candida auris*. Global phylogeny of clade III of *Candida auris* (n=566). The phylogeny was constructed from 2,152 SNPs. The color of the branches indicates the subgroup (referred as SG label) of each sample. The first ring (from inner to outer) indicates the isolate source (clinical or hospital-based environmental source), the second ring shows the country of origin of each isolate, the third ring indicates mutations related to fluconazole resistance, the fourth ring shows mutations related to micafungin resistance, and the last ring shows the year of culture of each isolate. Blank spaces in resistance rings mean that the isolates harbor the wildtype allele. Bootstrap support values of the 3 subgroups and the main outbreak/clusters are displayed.

Within subgroup SG3 (mean genetic distances 26 SNPs, ranging from 0-60 SNPs), there were two major clusters, one exclusive to the United States with 191 cases (189/191 isolates from California), and another, with a wider distribution, which included 372 samples from nosocomial outbreaks in Spain, UK, China and Kenya, as well as two small clusters from the United States with most cases from Florida (**Figure 2**). The mean genetic distance between isolates from the US cluster (only California cases) was 9 SNPs, while those of the outbreaks from Spain, UK, China, and Kenya were 2, 3, 6, and 10 mutations, respectively, confirming recent transmission events (**Figure 3**).

**Figure 2.**
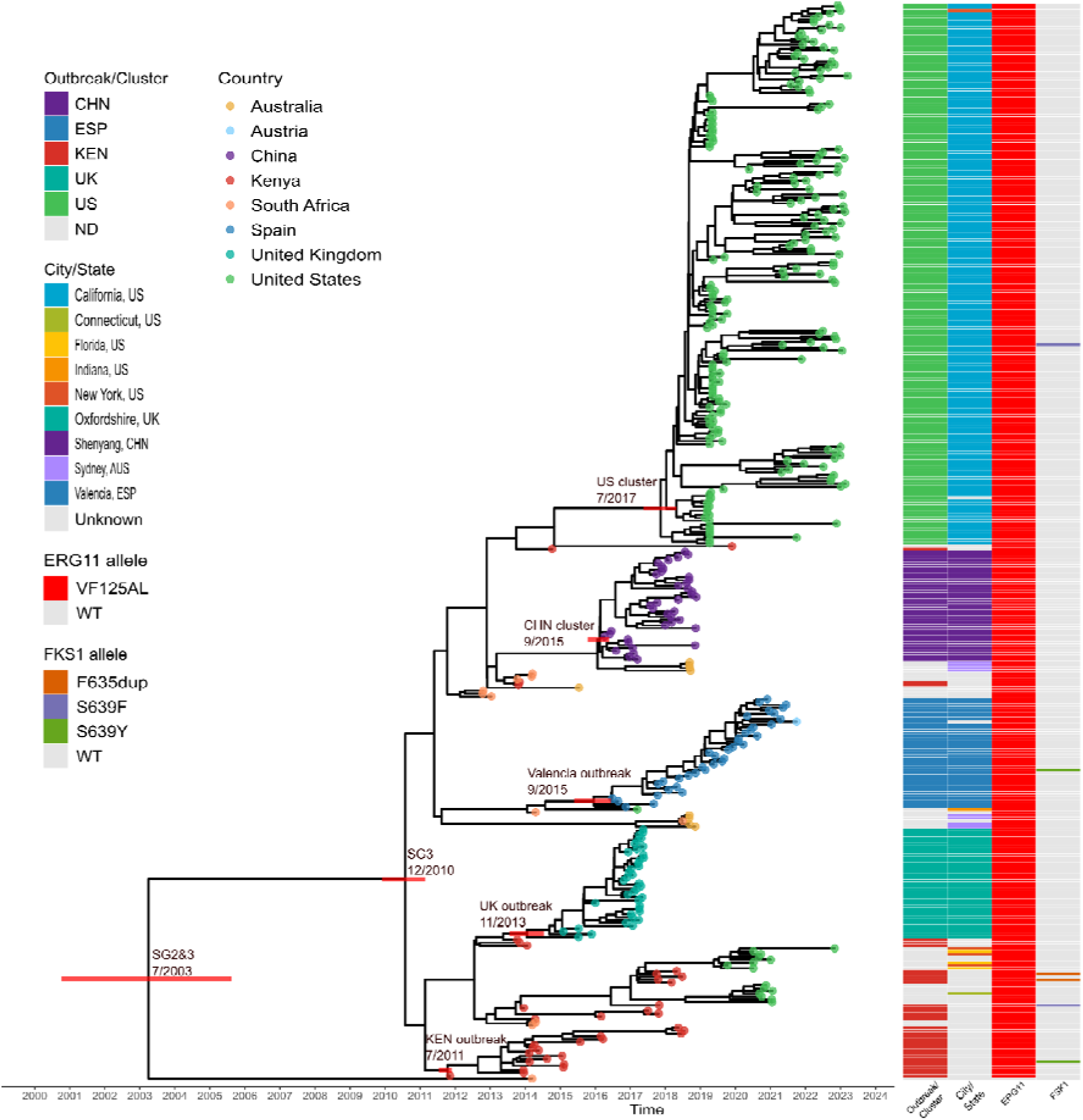
Dated phylogeny of *Candida auris* clade III subgroups SG2 & SG3 (n=380). The Bayesian-based tree was built from 1,071 SNPs. The inferred date (MM/YYYY) of the ancestral origin of SG3 and SG2&3, as well as that of the most recent common ancestor (MRCA) of the main clusters in this analysis are indicated at the corresponding nodes. Red bars indicate 95% HPD. Branch tip colors indicate the origin country of each sample analyzed. The first column indicates the outbreak or cluster that corresponds to each sample analyzed. The second column shows the city/state for samples with that information. The third column indicates mutations associated with fluconazole resistance (ERG11 allele), while the last column indicates mutations associated with micafungin (FKS1 allele). ND means "Not Detected".

**Figure 3.**
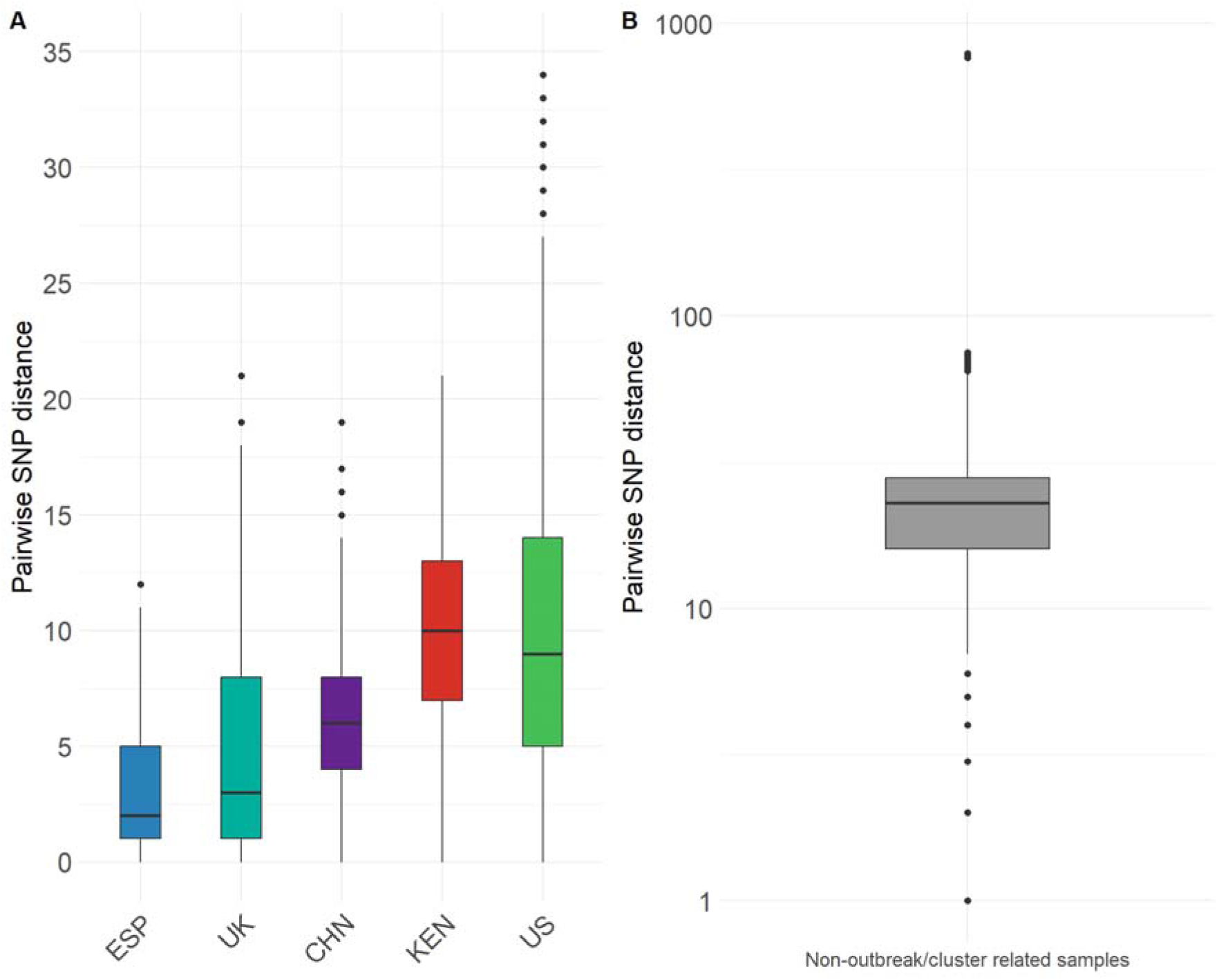
Pairwise genetic distances of *Candida auris* clade III subgroups SG2 & SG3. **A)** Genetic distances between isolates of each outbreak/cluster. **B)** Genetic distances of between samples not described as an outbreak or cluster.

Notably, we observed that almost all the isolates belonging to SG3 (except strain "B19006") carry the ERG11:VF125AL mutation associated with fluconazole resistance. In contrast, isolates classified in SG1 and SG2 possess the wildtype allele and are phenotypically susceptible to fluconazole. As for micafungin resistance, only 9 isolates contained a micafungin-related mutation. The most common mutation was FKS1:S639P with 3 cases, followed by FKS1:S639F, FK*S*1:S639Y, and FK*S*1:S639dup with two cases each.

### Local and global spread of clade III

Given the closer phylogenetic relationship between subgroups SG2 and SG3, we decided to focus our analysis on these subgroups and to examine their global dispersion. By considering the phylogenetic structure obtained through Bayesian analysis and the origin of each isolate, we found that SG3 originated in Africa, resulting in clinical cases in Kenya and South Africa between 2011 and 2014 (**Figure 2**).

Since then, cases have been reported from other continents. First in Oceania (Australia) and then in Europe (United Kingdom) in 2015 (20). A year later, in 2016, the first cases of nosocomial infections were reported in China and Spain (5, 21). In 2017, the first case was reported in the Americas: a patient in the United States in Indiana (strain "B12631") (10). However, this case may be considered as a single case, given that it did not lead to further noticed transmissions. It was not until 2019 when multiple cases of infection were reported in the states of California and Florida, the latter causing large nosocomial outbreaks in those locations (31). Finally, in 2020, isolated cases were described in Connecticut and New York. Importantly, since the first cases were reported in each locality, the pathogen continues to be identified. For instance, in Australia, new cases were reported in 2018, three years after the first described case (32).

### Dating analyses of clusters and outbreaks

Next, we analyzed subgroups SG2 and SG3 to infer the probable dates of origin of both lineages. For these analyses we used a dataset with samples having at least year/month in their available diagnosis date (n=380, MSA consisted of 1,071 SNPs) (see Methods). Once we ascertained that we had a strong temporal signal (R^2^=0.69) (**Supplementary Figure S2**), we decided to calculate the rate of evolution of both subgroups, as well as the most likely date of their most recent common ancestor (tMRCA). The inferred evolution rate was 2.25e-7 s/s/y (95% CI, 1.92e-7 - 2.46e-7) which translates as 2.87 changes per year (95% CI, 2.45 - 3.13). The tMRCA of the SG2 and SG3 subgroups was inferred in July 2003 (95% HDP, March 2001 – October 2005), while the origin of the fluconazole-resistant SG3 subgroup was December 2010 (95% HDP, April 2010 – June 2011) (**Supplementary Figure S3**).

Additionally, we obtained the most likely date for the MRCA of the nosocomial outbreaks as well as that of the US cluster. The dates of the Spain, UK, China, and Kenya clusters were September 2015 (95% HDP, February 2015 – March 2016), November 2013 (95% HDP, May 2013 – April 2014), September 2015 (95% HDP, April 2015 – January 2016), and July 2011 (95% HDP, April 2011 – August 2011), respectively, coinciding in all cases with the earliest reported cases in each corresponding setting (**Supplementary Figure S3**). The ancestor of the large US cluster likely originated in July 2017 (95% HDP, February 2017 – May 2018). The states where this subgroup has been reported are California with 189 cases, followed by Florida with 75 cases, New York with four isolates, and Connecticut and Indiana with one case each. Looking at the phylogeny of clade III we may suggest that there were multiple introductions in the US, one in New York and one in California. The latter is the cause of a large outbreak since 2017. This geographic restriction suggests that the clades have diversified locally after their ancestral strains were introduced into the region (**Supplementary Figure S4**).

## DISCUSSION

In this work we have analyzed the evolution of clade III of *Candida auris* and described how it has spread and established in the world causing nosocomial, antifungicide-resistant outbreaks and clusters in all the continents. To the best of our knowledge, this is the first study in which the evolution of this clade is examined in detail. We found that this clade is divided into three subgroups (SG1, SG2 and SG3). Of these, subgroups SG2 and SG3 are genetically more closely related, while SG1 is quite distinct at the genetic level.

The strains that belong to subgroup SG3 are resistant to fluconazole and carry the VF125AL mutation in the ERG11 protein. As a result, the use of other antifungal drugs is essential; thus, the emergence of multidrug-resistant isolates is highly probable. However, the frequency of known resistance-related mutations to these antifungals is low. We observed only 9 cases (1.6% of the total) with mutations in the FKS1 gene associated with resistance to micafungin. This might be due to the fact that there are unknown mutations associated with these compounds, or that many isolates lack this phenotypic information, or that there are other mechanisms of resistance that remain unknown. As a practical case in this study, strain "B19006" does not carry mutation VF125AL:ERG11. This isolate probably carries other variants (SNPs, deletions or insertions), or alterations in the transcription of unknown genes that may be associated with resistance. Recently, new mutations or target genes related to resistance have been described (33).

Based on the ancestral dates inferred from the evolutionary analysis, we were able to determine the origin, timing and introduction of clade III at each setting. By using available genomes and the dates of different previously reported outbreaks, we have established that the likely origin of SG3 was in Africa in December 2010 (95% confidence interval: April 2010 - June 2011), from which an outbreak originated in a facility in Kenya in mid-2011 (July 2011) (34). In 2013, a major outbreak was reported in a hospital in the United Kingdom (20), coinciding with our obtained estimates.

In the second half of 2015, the first reported cases were described simultaneously in a hospital in the Shenyang province, China, as well as in Valencia, Spain. Both dates coincide with those reported in both areas (5, 17). Finally, the SG3 subgroup arrived in the United States in 2017 (July 2017) through two introductions, one in New York and another in California. The New York isolates are more closely related to isolates described in South Africa, suggesting an African ancestry, while those from California are more restricted to that area and their most recent ancestry is unknown. The first cases reported in California are from late 2017 (31).

Fluconazole resistance and the origin of clade III have been widely debated. In this study, we identified two genetically closely related subgroups (SG2 and SG3), one susceptible and the other surprisingly resistant to the aforementioned antifungal drug. Considering the inferred dates of the possible ancestry of both subgroups and SG3 (7/2003 and 12/2010, respectively), we suggest that the acquisition of resistance in SG3 occurred during this period. We hypothesize that the increased use of antifungals in both clinical and other ecological niches has exerted selective pressure for the emergence and fixation of resistant strains. In fact, increased use of antimicrobials has been reported in South Africa between 2004-2006 (3). This together with the fact that in South Africa there are many reported cases of fungal infections caused by other *Candida* and non-*Candida* species, which have been treated with fluconazole (35), suggests that the likely origin of this resistant phylogenetic subgroup was in South Africa and extended worldwide.

A noticeable result is the estimated evolutionary rate. In this study the estimated mutation rate as 2.25e-7 s/s/y (95% HPD, 1.92e-7 - 2.46e-7), while the mean rate reported in other studies has an average value of 3.02e-5 s/s/y (4, 14, 15). This discrepancy may be due to different factors regarding how this value was obtained. The main difference is the number of samples involved. In our study we used 380 complete genomes from clade III, aimed at including as much diversity as possible, while in other studies smaller datasets were used, mainly genomes from a transmission outbreak to calibrate estimates for the whole clade. Since more genomes are now available, the inferred rate of evolution is more accurate and likely closer to reality. Furthermore, the R^2^ value we obtained is higher than in other studies (0.69 vs. 0.35 and 0.55), giving a higher confidence to the inferred rate. Another important difference that might affect the evolutionary rate estimate is the reference genome used. In other studies, the reference genome of clade I (strain B8441, GCA_003014415.1) was used, whereas in our analysis we selected the reference genome of clade III (strain B12221, GCF_002775015.1). The impact of the genome used when performing this kind of studies has been discussed elsewhere, from the percentage of reads mapped to the number of variants called (36). Considering all these points, we consider our approach to be more accurate and robust (4, 10).

The limitations of this study are based on the number of samples analyzed. Although many epidemiological studies report isolates belonging to clade III (4, 10, 18, 37), there are other clades that are more successful, such as clade IV. We suggest that this may be due to the fact that the latter arrived first and rapidly colonized these regions (4).

In conclusion, we have analyzed complete genomes of *Candida auris* clade III to provide new insights into the origin of a fluconazole-resistant subgroup, and its dispersal dynamics over the world. These results open new discussions and extend the knowledge to implement epidemiological measures to control the pathogen, especially in healthcare settings.

## ACKNOWLEDGMENTS

We would like to thank all the research groups, medical units, hospitals and especially the surveillance systems for generating, depositing and making available all the genomic data used in this study. Unpublished genomic data downloaded from NCBI were deposited therein by the U.S Centers for Disease Control and the National Center for Emerging and Zoonotic Infectious Diseases-Mycotic Diseases Branch.

Irving Cancino-Muñoz was supported by a Margarita Salas contract (UNI/551/2021) from Ministerio de Universidades (Spanish Government). This work was supported by projects CIPROM-2021-053 from Generalitat Valenciana and PID2021-127010OB-I00 from Ministerio de Ciencia e Investigación (Spanish Government).

## SUPPLEMENTARY FIGURES

**Supplementary Figure S1.**
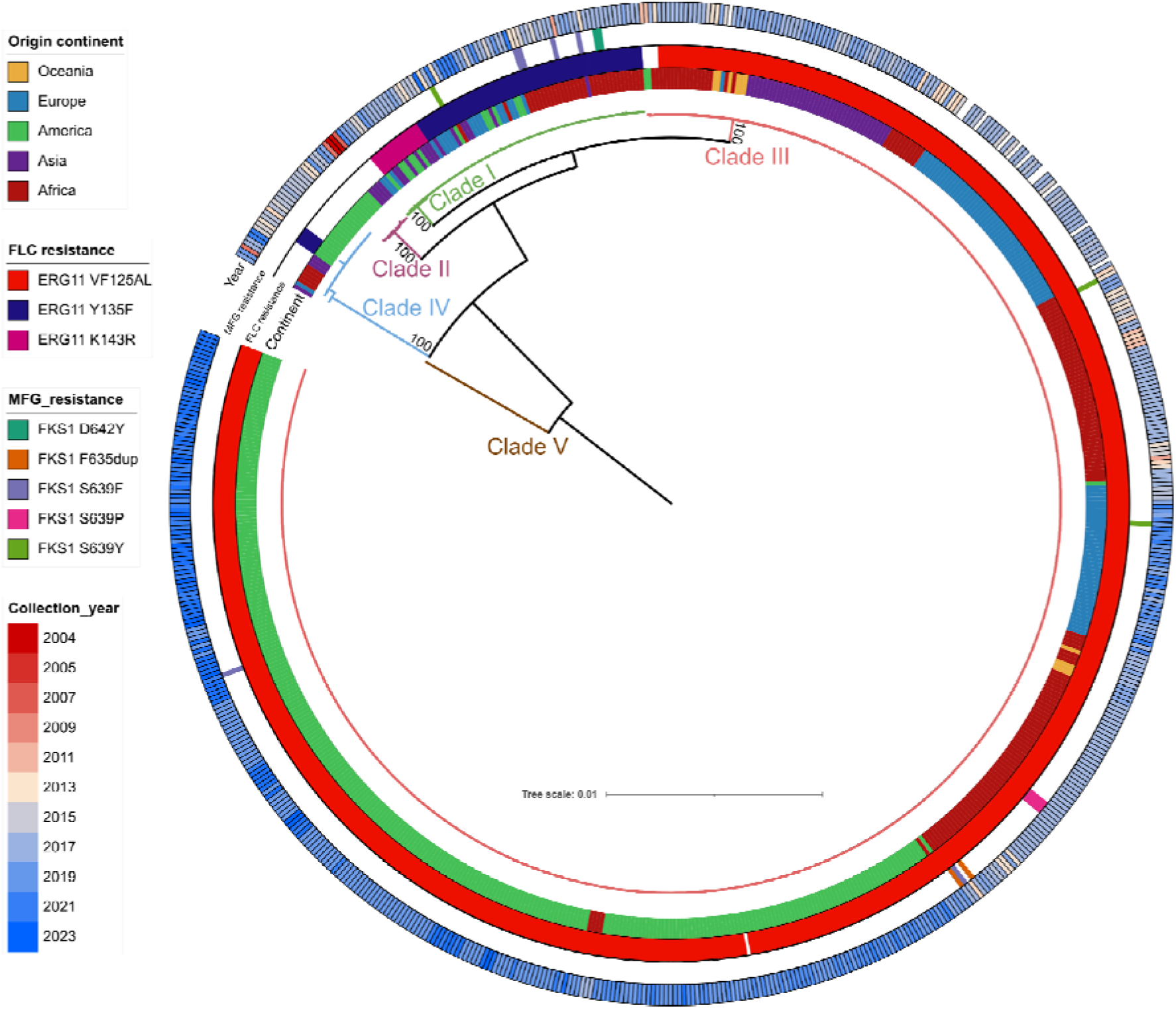
Global phylogeny of all *Candida auris* isolates (n=689) obtained by maximum likelihood. The phylogeny was constructed from 194,389 SNPs. The color of each sample indicates the continent of origin. The first ring (from inner to outer) shows the sample origin, the second ring indicates mutations related to fluconazole resistance, the third ring shows mutations related to micafungin resistance, and the last ring shows the year of culture of each isolate. Blank spaces indicate that the isolates harbor the wildtype allele. Bootstrap support values of all *C.auris* clades are displayed.

**Supplementary Figure S2.**
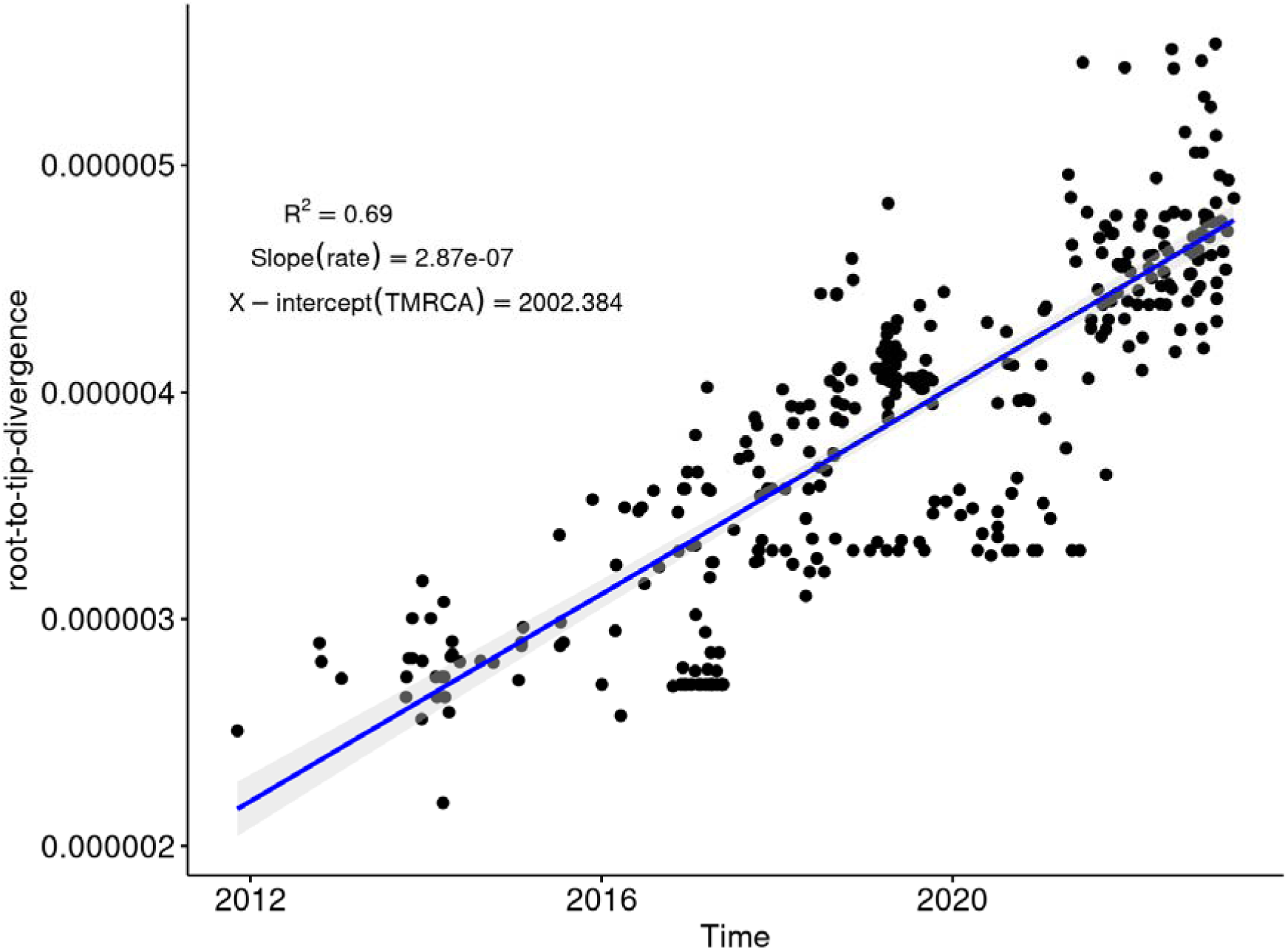
Root-to-tip regression analysis of *Candida auris* clade III genomes analyzed. The slope represents the evolutionary rate obtained in Tempest. Each point represents a different genome. Shadow area around the regression line represents its 95% CI.

**Supplementary Figure S3.**
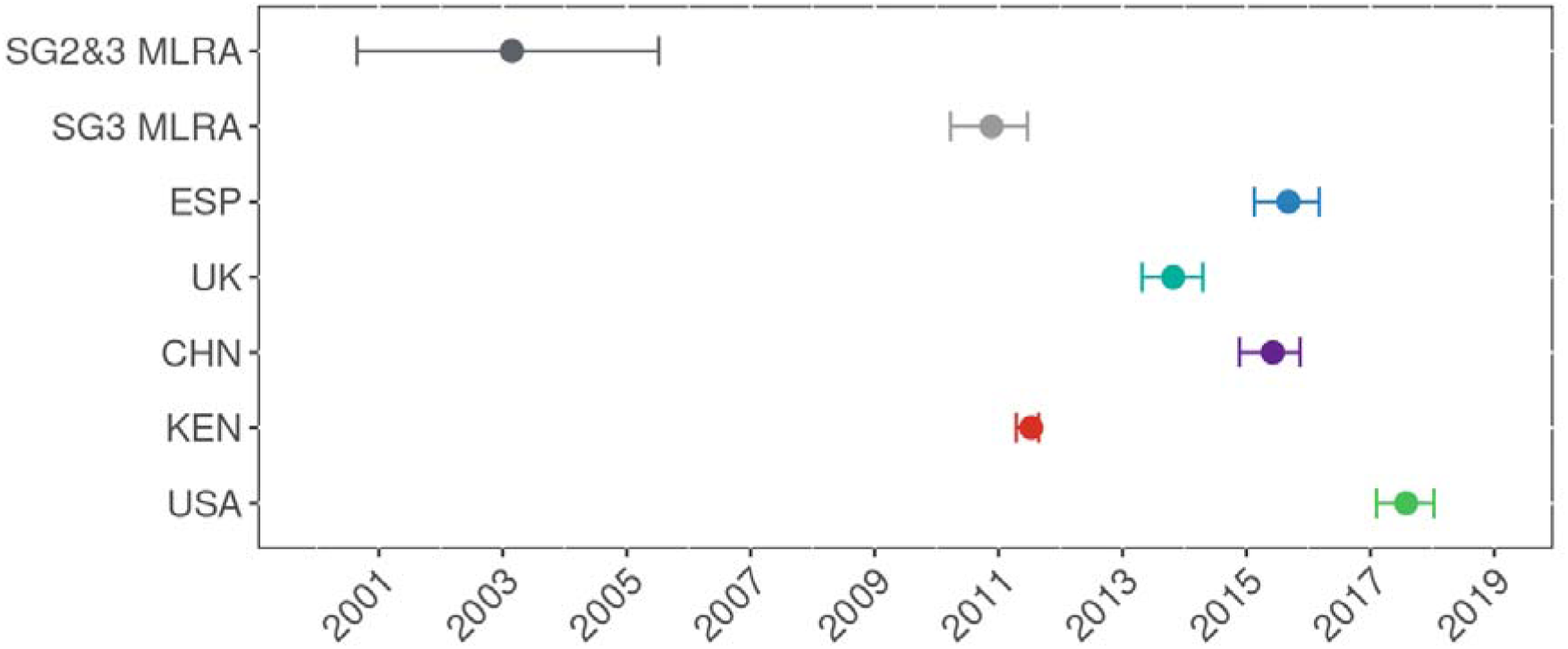
Inferred dates of the MRCA of each outbreak/cluster analyzed and of those of SG2&3 and SG3, the fluconazole-resistant subgroup. Confidence intervals for each inferred point (95% CI) are shown in the graph.

**Supplementary Figure S4.**
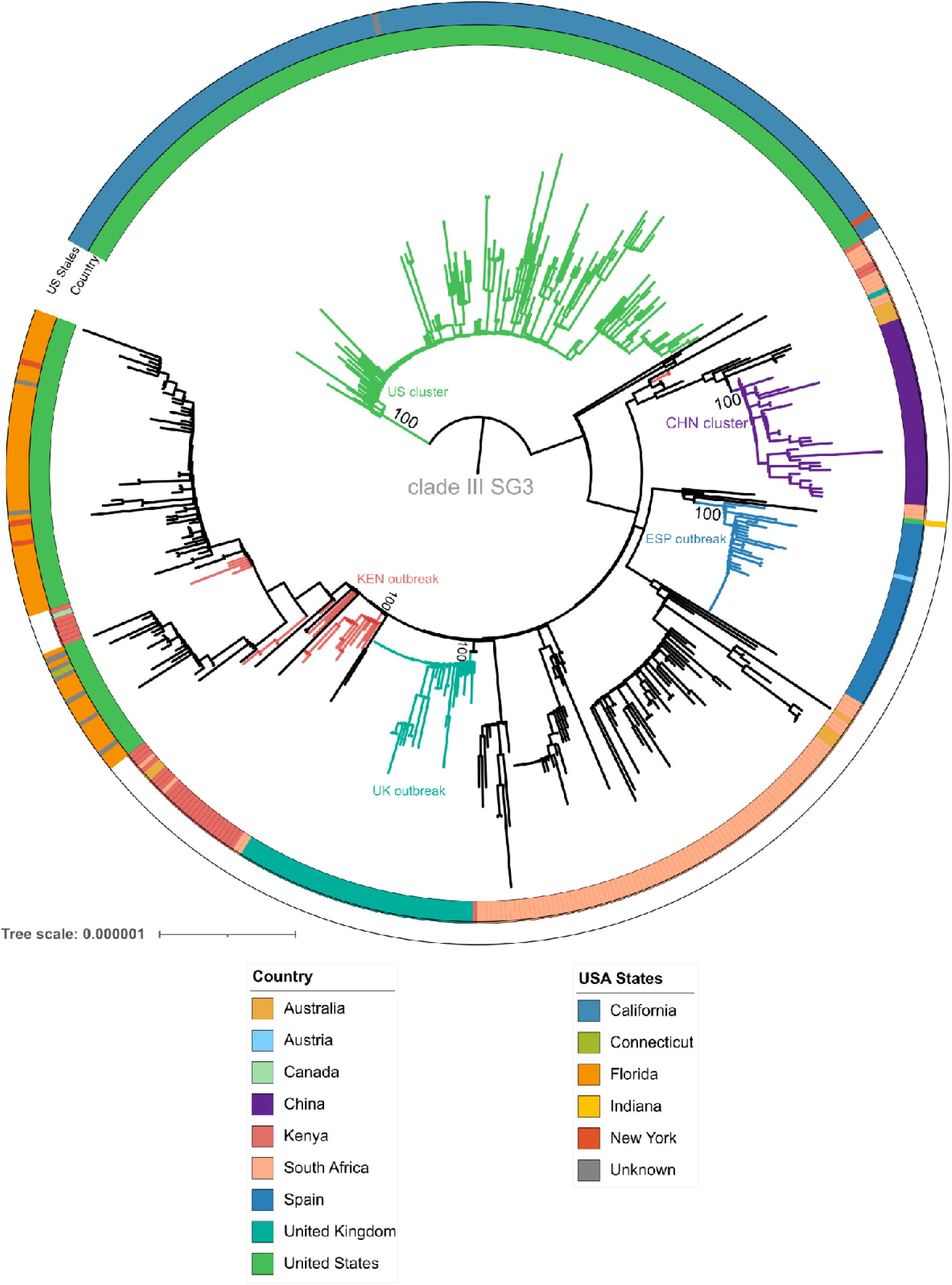
Global phylogeny of all SG3 *Candida auris* isolates analyzed (n=562). The first ring (from inner to outer) indicates the origin country of each sample, the second ring shows the corresponding US state of the USA samples. Branch colors represent the different outbreaks analyzed as well as the US cluster defined in the study. Bootstrap support values of all outbreak/clusters are displayed.

## Supplementary Tables Legends

**Supplementary Table 1.** Sequencing data of all *Candida auris* genomes analyzed in this study (n=689).

**Supplementary Table 2.** Metadata (sample origin, phenotypic and genotypic characteristics of resistance-related mutations) of all *Candida auris* analyzed in this study (n=689).

**Supplementary Table 3.** Metadata (sample origin, phenotypic and genotypic characteristics of resistance-related mutations) of all *Candida auris* Clade III genomes analyzed in this study (n=566).

**Supplementary Table 4.** Metadata (sample origin, phenotypic and genotypic characteristics of resistance-related mutations) of all *Candida auris* Clade III genomes used for dating analysis (n=380).

